# Intentional stocking undermines ecological stability

**DOI:** 10.1101/2022.02.24.481650

**Authors:** Akira Terui, Hirokazu Urabe, Masayuki Senzaki, Bungo Nishizawa

## Abstract

The past decades have witnessed efforts to unveil ecological risks associated with massive releases of captive-bred individuals (“stock enhancement”). Still, we may underestimate the negative impact because current schemes rarely consider the consequences of stock enhancement for the whole ecological community. Here, we use theory and long-term data of Japanese stream fish communities to show that stock enhancement undermines community stability. Our theory predicted that stock enhancement destabilizes community dynamics by facilitating competitive exclusion. Consistent with this prediction, fish communities showed greater temporal fluctuations and fewer species richness in rivers with the intensive stocking of hatchery masu salmon - a major freshwater resource in the region. Our findings paint a bleak picture for stock enhancement, reinforcing the recurrent calls for alternative management strategies.

**One Sentence Summary:** Large-scale releases of captive-bred organisms undermine the long-term persistence of ecological communities by disrupting the stable coexistence of competing species.

## Maintext

Human demands for natural resources are ever-increasing, such that active interventions are critical to the sustainable management of fisheries, forestry, and wildlife (*1*). Captive breeding is a form of the efforts to enhance wild populations of diverse plant and animal taxa (*1, 2*). Although releases of captive-bred organisms (“stock enhancement”) entail ecological risks such as the accumulation of deleterious alleles (*3, 4*), this method is still pervasive in conservation (*2*) and natural resource management (*1*). In fisheries, for example, billions of hatchery individuals are released annually across the globe (*5*). The widespread use of captive breeding is perhaps because of the “myth” that the demographic benefits of released individuals may exceed the associated risks.

Current debates, however, overlook the fact that we have rarely assessed the community-wide impact of stock enhancement. Species are all embedded in the complex web of interacting organisms, and the stable coexistence of competing species through density-dependent feedback underpins the emergent stability of ecological communities (*6–8*). Stock enhancement may disrupt the sensitive balance of species interactions because it introduces unnaturally high numbers of individuals into the wild (*9*). Hence, this form of species management may intervene in the ecological process that allows competing species to coexist, ultimately degrading the long-term community stability. Evidence for this hypothesis is lacking, however.

Here, we show that stock enhancement undermines long-term community stability, which we define as the relative size of fluctuations in total community density over time (*6*). Our theory illuminates that stock enhancement compromises the stabilizing mechanism emerging from species niche differences. The present study further demonstrates the relevance of this general theory to natural systems by showing its congruence with Japanese stream fish communities, where ∼10 million hatchery masu salmon (*Oncorhynchus masou masou*) are released annually for fisheries, recreation, and conservation purposes (*10*).

We employed a multispecies Ricker model (*11*) to simulate community dynamics with the selective stock enhancement of a constituent species (species 1). Specifically, the population density of species *i* at time *t* + 1, *N*_*i,t*+1_, is modeled as:

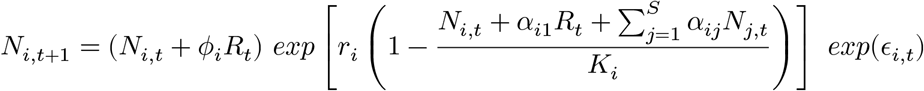

where *r*_*i*_ is the intrinsic growth rate, *α*_*ij*_ the competition coefficient of species *j* on species *i, K*_*i*_ the carrying capacity, *R*_*t*_ the number of released individuals for the enhancement of species 1, and *ϵ*_*i,t*_ the species response to stochastic environmental fluctuations that follow a normal distribution 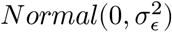. To capture variation in species traits, intrinsic growth rates of unenhanced species and interspecific competition coefficients were drawn randomly from a uniform (*r*_*i,i*≠1_ ∼ *Unif*(0.5, 2)) and an exponential distribution 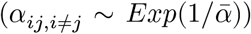. The parameter *ϕ*_*i*_ controls the relative fitness of captive-bred individuals as follows:

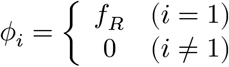

*f*_*R*_ (≥ 0) is the density-independent survival of captive-bred individuals relative to wild individuals. Therefore, the model accounts for the fitness difference of captive-bred individuals due to genetic effects and/or plasticity (*2, 12, 13*) when considering the reproductive contribution to the next generation. Without loss of generality, we assumed constant *K*_*i*_ (*K*_*i*_ = *K*) and *R*_*t*_ (*R*_*t*_ = *R*) across species and time, respectively.

We ran 1600 time steps of 1000 independent communities (i.e., simulation replicates) under each of 24 simulation scenarios. These scenarios cover a range of ecological contexts, differing in the intrinsic growth of an enhanced species *r*_1_, competition 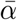, carrying capacity *K*, and relative fitness *f*_*R*_ (see **Materials and Methods**). Using the last 1000 time steps, we obtained the following summary statistics of the total community density 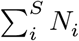 to examine the community-level response to stock enhancement: coefficient of variation (CV), number of species persist (defined as *N*_*i*_ > 0.01 at *t* = 1600), temporal mean (*μ*), and temporal SD (*σ*). We also calculated the temporal mean and SD for the enhanced (*N*_1_) and unenhanced species 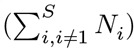 separately to infer underlying mechanisms.

When carrying capacity is small (*K* = 100), our model predicted a destabilizing effect of stock enhancement on ecological communities as illustrated by increased CV with increasing numbers of releases (**Figure 1A**). This pattern stemmed mainly from the reduced mean of the total community density, and both enhanced and unenhanced species groups were responsible (**Figure 1A**). The enhanced species decreased because stocking induced the negative competitive effect exceeding the reproductive contribution of released individuals (*14*). Meanwhile, interspecific competition reduced the unenhanced species at high levels of stock enhancement, resulting in fewer persisting species (**Figure 1A**). Combined, the total community density decreased more sharply than individual species groups (**Figure 1A**). The SDs showed a similar trend, but the relationship was flatter at the community level (**Figure 1A**). Since a CV is a ratio of an SD to a mean, the steeper decline of the mean community density led to the increased CV. These patterns were qualitatively similar across most ecological contexts (robust to the changes in *r*_1_, 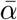, and *f*_*R*_; **Figures S1-S3**).

**Figure 1.**
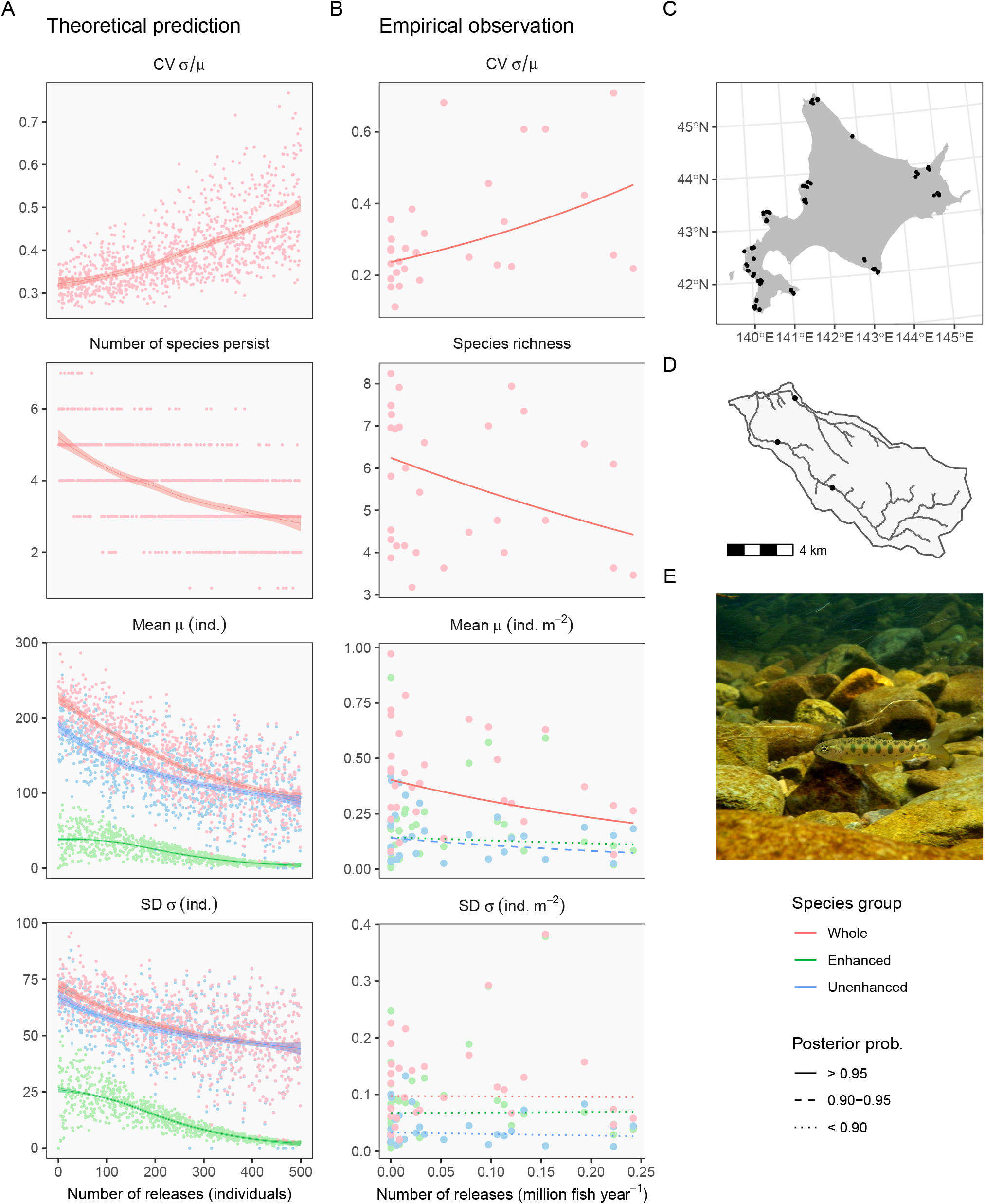
Theory and empirical observations agree that stock enhancement destabilizes community dynamics. In A and B, panels and colors distinguish response variables and species groups. (A) Theoretical predictions. Dots represent individual simulation replicates, and lines and shades are loess curves fitted to simulated data and 95% confidence intervals. Parameters used in this simulation are: intrinsic growth rate of an enhanced species *r*_1_ = 1, average interspecific competition 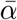 = 0.5, carrying capacity *K* = 100, environmental stochasticity *σ*_*ϵ*_ = 0.5, relative fitness of captive-bred individuals *f*_*R*_ = 0.5. (B) Empirical evidence. Dots represent geometric means of site-level observations in 31 watersheds. Lines are the predicted values of the regression models, and line types correspond to the coefficient’s posterior probabilities. See Tables S8-10 for full statistics. (C) Map of sampling sites (black dots) in Hokkaido, Japan. (D) Example of a protected watershed (Okushibetsu). (E) Masu salmon *Oncorhynchus masou masou*. Photo credit: Akira Terui.

The destabilizing effect emerges because stock enhancement affects the balance of species interactions that underpins community stability. In theory, the stable coexistence requires a niche difference that is large enough to overcome the relative difference in intrinsic competitive ability (*7*). Under this condition, competing species can grow from small populations because dominant species undergo stronger intraspecific competition (*7*). Such coexistence favors stable temporal dynamics of species-rich communities (*6*) because it gives rise to “overyielding” (*7, 8*), i.e., total community density of a multispecies community exceeds what would be expected in a single species community 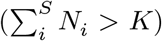. However, stock enhancement is externally controlled, and the number of releases is not subject to density-dependent regulation. Therefore, released individuals impose additional intra- and interspecific competition that interferes with the ecological process producing overyielding.

Stock enhancement, however, had little influence on community dynamics when carrying capacity was sufficiently large (*K* = 500; **Figures S4-S6**). In particular, stock enhancement increased the enhanced species with a low population growth rate (*r*_1_ = 0.5; **Figure S4**). This result may explain why some of the best evidence for successful stock enhancement comes from long-lived endangered species (e.g., *15*). More importantly, the contrasting community response at different carrying capacities provides deeper insights. One of the core motivations for stock enhancement is to mitigate declining trends of natural resources due to human impacts, such as habitat loss (*16*). Ironically, our results suggest that stock enhancement will not bring desired outcomes unless we resolve root causes that compromise environmental capacity first.

To demonstrate the relevance of our general theory to natural systems, we assessed the potential impacts of the stock enhancement of masu salmon (**Figure 1E**) on the long-term stability of stream fish communities in Hokkaido, Japan. In the protected watersheds (all separated by the ocean; **Figure 1C, D**), a long-term program exists to monitor stream fish communities along with the official stocking records. The majority of stocking occurs in spring, after which salmon fry stay in freshwater for growing seasons. Therefore, the study system sets the stage for a “natural experiment” to test our theoretical predictions. We used the data from 1999 to 2019 at 97 sites within 31 independent watersheds (see **Materials and Methods** for selection criteria). Using hierarchical Bayesian models, we quantified the effect of stock enhancement on community dynamics while accounting for potential effects of climates and local abiotic factors.

As predicted, stream fish communities showed greater temporal fluctuations (higher CV) in watersheds with intensive stocking (**Figure 1B**). The effect was striking in its magnitude, almost doubling the CV at the highest stocking level. Our analysis strongly supported the positive relationship between the CV and the number of releases (**Figure 2**), in which the probability of the regression coefficient being positive was 0.99 (**Table S8**). This pattern was associated with the reduced long-term average of the total community density and fewer species richness (**Figures 1B and 2**), and both enhanced (masu salmon) and unenhanced fish groups contributed to this trend (**Figures 1B and 2;** see **Tables S8-S10** for full statistics). In the meantime, the SDs had vague relationships with stock enhancement (**Figures 1B and 2**).

**Figure 2.**
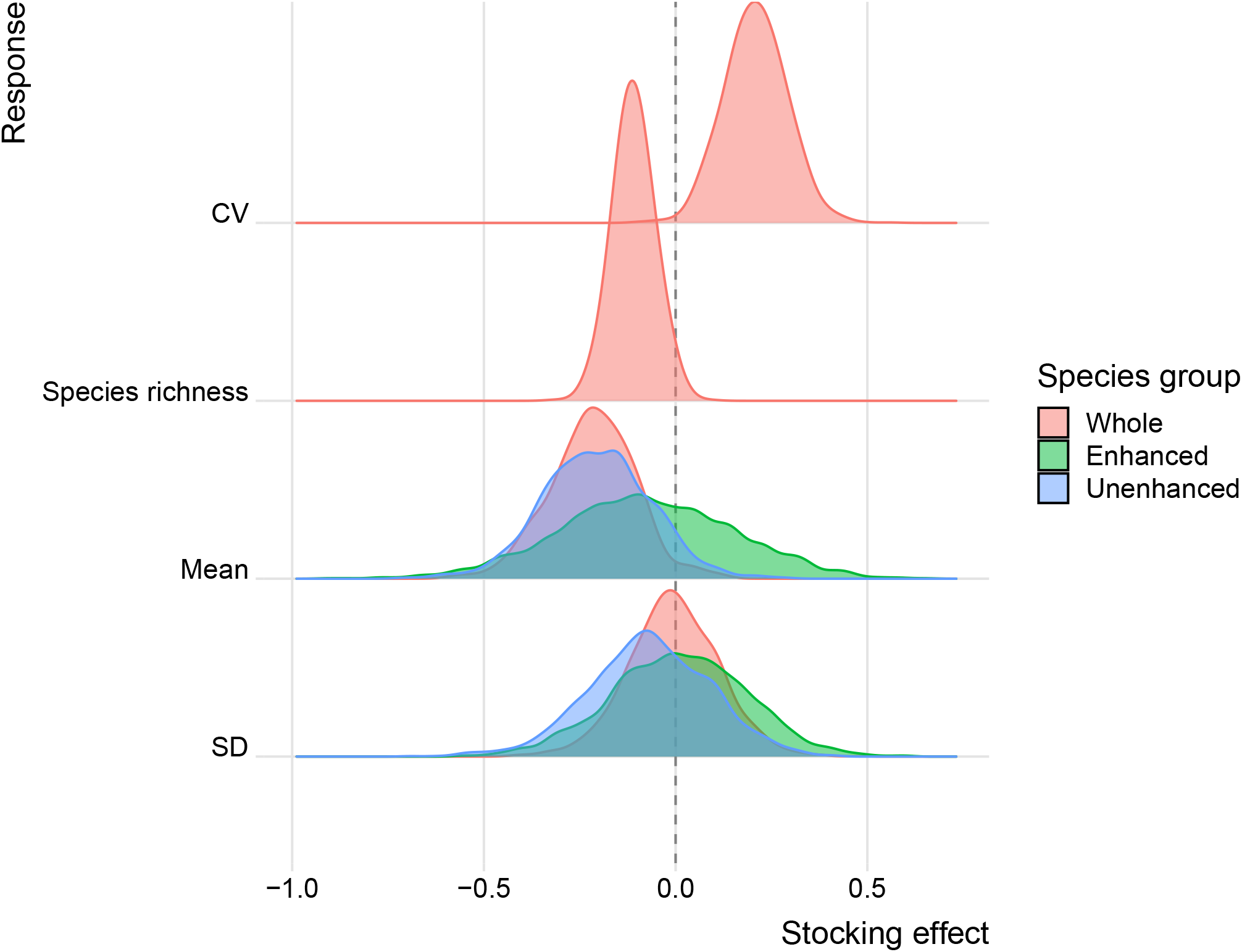
Posterior distributions for the standardized regression coefficients of stock enhancement. Y-axis represents different response variables grouped by colors distinguishing species groups (whole community, enhanced species [masu salmon], and unenhanced species).

Competition is a likely mechanism, as our theory assumes. Hatchery salmons are larger and more aggressive than wild individuals, increasing the likelihood of intense intra- and interspecific competition in the wild (*12*). Indeed, field and experimental studies confirmed that hatchery masu salmon competed with wild masu salmon and other stream fishes during their freshwater life stage, typically one year (*17, 18*). Importantly, socioeconomic factors control the number of releases (e.g., budget allocation) regardless of the current condition of recipient communities. As such, released fish are probably “excessive” and may cause resource competition that would otherwise not exist.

The reduced fitness of hatchery masu salmon may also play a role in the observed response. Like other salmonids, ocean-migrating adults of hatchery masu salmon show lower return rates to the spawning river (*19*). In addition, the aggressive behavior of hatchery salmons may make them vulnerable to predation (*12*). The fitness disadvantage may add to negative density dependence to influence community stability.

We cannot exclude the possibility that spurious correlations drove our results. However, the protected watersheds preserve nearly intact landscapes (**Figure S10**) with strict regulations of human activities (e.g., exploitation, construction of in-stream structures); therefore, it is difficult to envision that unmeasured human influences caused the observed relationships. Further, we have statistically controlled natural variation in environmental factors (see **Materials and Methods** and **Tables S8-S10**). This unique setup may have helped uncover the qualitative agreement between theory and empirical patterns.

Despite the significant attention to the fate of captive-bred individuals (*2, 13*), current schemes rarely consider the self-regulation process of biodiversity. Our results suggest that the ignorance of this critical process may erode the long-term persistence of the recipient community, likely impacting the provisioning of ecosystem services (*20*). While our empirical example is limited to stream fish communities, we anticipate that this phenomenon is pervasive in nature because the destabilizing effect emerged across diverse simulated scenarios.

We should prioritize habitat conservation with a broader scope of ecosystem management. Such efforts are particularly important in the Anthropocene because habitat degradation may exacerbate the undesired influence of stock enhancement. Protected areas and environmental restoration are promising tools to conserve biodiversity, and a smart spatial design is a key to achieving successful conservation. For example, coordinated placement of conservation sites considering spatial biodiversity patterns is crucial in improving the ecological outcomes (*21–23*). Governance may also play a central role in enforcing environmental legislation, potentially determining the effectiveness of conservation investment (*24*). These considerable potentials indicate that viable management options exist before blindly accepting stock enhancement. Without a comprehensive framework that appreciates the ecological integrity of natural communities, the stock enhancement will never be effective but impairs biodiversity.

## Acknowledgments

We are grateful to people involved in the long-term monitoring program at the protected watersheds in Hokkaido. We thank Masato Yamamichi for helpful discussion on this manuscript. Data and codes will be made available upon acceptance of the manuscript.

