## Supplementary Materials for "Intentional stocking undermines ecological stability"

**This PDF file includes:**

- Materials and Methods
- Supplementary text
- Tables S1-10
- Figures S1-S11
- References

**References** **27**

### Materials and Methods

#### Theory

We employed a multispecies Ricker model (*1*). In the basic formula without stock enhancement, the population density of species $i$ at time $t+1$, $N_{i,t+1}$, is modeled as:

$$N_{i,t+1} = N_{i,t}\ exp\left[r_i\left(1 - \frac{N_{i,t} + \sum_{j=1}^{S}\alpha_{ij}N_{j,t}}{K_i}\right)\right] exp(\epsilon_{i,t}) \quad (1)$$

where $r_i$ is the intrinsic growth rate, $\alpha_{ij}$ the competition coefficient of species $j$ on $i$, $K_i$ the carrying capacity, and $\epsilon_{i,t}$ the species response to stochastic environmental fluctuations that obey an normal distribution $Normal(0, \sigma_\epsilon^2)$. We modified this formula to include the effects of stock enhancement (species 1) on reproduction and competition as follows:

$$N_{i,t+1} = (N_{i,t} + \phi_i R_t) exp\left[r_i\left\{1 - \frac{N_{i,t} + \alpha_{i1}(N_{1,t} + R_t) + \sum_{j=2}^{S}\alpha_{ij}N_{j,t}}{K_i}\right\}\right] exp(\epsilon_{i,t}) \quad (2)$$

$R_t$ is the number of released individuals, and the parameter $\phi_i$ controls the relative fitness of captive-bred individuals:

$$\phi_i = \begin{cases} f_R & (i = 1) \\ 0 & (i \neq 1) \end{cases} \quad (3)$$

$f_R$ ($\geq 0$) is the density-independent survival of captive-bred individuals relative to wild individuals. Equation (2) can be reorganized to:

$$N_{i,t+1} = (N_{i,t} + \phi_i R_t) exp\left[r_i\left(1 - \frac{N_{i,t} + \alpha_{i1}R_t + \sum_{j=1}^{S}\alpha_{ij}N_{j,t}}{K_i}\right)\right] exp(\epsilon_{i,t}) \quad (4)$$

In this model, intrinsic growth rates of unenhanced species $r_{i,i\neq1}$ and interspecific competition $\alpha_{ij}$ are random draws from a uniform ($r_{i,i\neq1} \sim Unif(0.5, r_{max})$) and an exponential distribution ($\alpha_{ij} \sim Exp(1/\bar{\alpha})$), respectively. We assumed constant values of intraspecific competition ($\alpha_{ii} = 1$), carrying capacity ($K_i = K$) and the number of releases ($R_t = R$).

Prior to the main simulation, we performed an extensive sensitivity analysis to identify parameters that strongly influence the relationship between community dynamics and stock enhancement (see **Supplementary text**). We identified five influential parameters (**Table S2-S4**), of which different values were considered in the main simulation as follows: intrinsic growth of enhanced species ($r_1$ = 0.5, 1, 2), average strength of interspecific competition ($\bar{\alpha}$ = 0.1, 0.5), carrying capacity ($K$ = 100, 500), and relative fitness of captive-bred individuals ($f_R$ = 0.5, 1). Meanwhile, we fixed values of the following parameters: the number of species ($S$ = 10), maximum intrinsic growth rate of unenhanced species ($r_{max}$ = 2), and environmental stochasticity ($\sigma_\epsilon$ = 0.5). This simulation setup resulted in 24 sets of parameter combinations that cover a range of ecological scenarios.

Under each parameter combination, we ran 1600 time steps of 1000 independent communities (i.e., simulation replicates). The number of released individuals $R$ was drawn randomly from a uniform distribution for each simulation replicate as $R \sim Unif(0, 500)$. We initialized the community with populations of each species drawn from a Poisson distribution with a mean of five. We repeated the seeding procedure every 10 time steps over the first 200 time steps to allow species to establish populations and reach equilibrium without stock enhancement (initialization period). After the initialization period, we released $R$ individuals of the enhanced species every time step over the next 400 time steps to reach new equilibrium with selective stock enhancement (burn-in period). We continued the simulation run with stock enhancement and saved the last 1000 time steps. We obtained the following summary statistics of the whole community $\sum_i^S N_j$: the coefficient of variation (CV), the number of species persist (defied as $N_i > 0.01$ at $t = 1600$), the temporal mean ($\mu$), and the SD ($\sigma$). The temporal mean and SD were also calculated for the enhanced species ($N_1$) and unenhanced species ($\sum_{i,i\neq 1}^S N_i$) separately. We summarized values of simulation parameters in **Table S1**.

#### Empirical analysis

##### Data

**Time-series data.** We assembled time-series fish data at 126 sites within 32 protected watersheds of Hokkaido Island, Japan. The Hokkaido Research Organization leads a long-term monitoring program at these watersheds, and the data are published as annual reports (*2*). The program began in 1963, but an effective, standardized sampling method has been implemented since 1999 (two-pass sampling with a combination of electrofishing and cast net). Most data were collected in summer with irregular interannual intervals (1- to 3-year intervals for most cases), and sampling efforts were quantified by sampling area (average: 175.49 ± 115.91 m$^2$). We confined our analysis to the sites that meet the following criteria: (i) the observation span (from the first to the last year of observation) exceeds 10 years, (ii) the number of observation years exceeds five years, and (iii) masu salmon is observed at least twice during the observation period. As a result, we used time-series data at 97 sites within 31 watersheds from 1999 to 2019. Summed abundance of first and second passes was used in the following analysis. **Table S5** summarizes observed species in these watersheds.

**Fish stocking.** The stocking of masu salmon began in the 1950s. Hatchery fish are released in spring (fry and smolt stages) and fall (juvenile stage). Although the fish release occurs at multiple locations within a watershed, the stocking information is available only at the watershed level. For each release stage, we assembled annual records of stock enhancement (the number of fish released; 1999-2019) from annual investigations by the Japan Fisheries Research and Education Agency and Salmon and Freshwater Fisheries Research Institute. During the study period, the majority of stocking took place in spring at a fry stage (fry : juvenile : smolt = 1 : 0.09 : 0.41)

**Environmental data.** At each sampling site, we measured the following environmental variables as potential covariates: upstream watershed area (km$^2$; a proxy for stream size), proportional land use in the upstream watershed (forest, urban, agriculture), local climates (annual mean air temperature [$^\circ C$] and cumulative precipitation [mm]), and ocean productivity (sea surface chlorophyll *a* concentration [mg m$^{-3}$]). We used MERIT Hydro (*3*) to delineate the upstream watershed polygon for each sampling site. We estimated the proportion of forest, urban, and agriculture in each watershed polygon based on land use data in 2015 from Copernicus Global Land Service (100-m resolution) (*4*). Climate data at each sampling site were extracted from CHELSA version 1.2 (*5*, *6*). We extracted annual data of chlorophyll *a* concentration (2002-2019; resolution, 4.6 km$^2$) from OceanColor (*7*) as a proxy for ocean productivity and calculated the average value within the 30-km radius of each river mouth. We used the following R packages to perform geospatial analysis: *sf* (*8*), *raster* (*9*), *exactextractr* (*10*), *stars* (*11*), *whitebox* (*12*).

##### Statistical analysis

Our goal is to compare temporal community dynamics across sites. However, the data are not comparable because of observation errors (e.g., different observers) and missing observations. To confront this challenge, we developed a Bayesian state-space model for three species groups separately: (i) whole community, the summed abundance of all species, (ii) enhanced species, the abundance of masu salmon, (iii) unenhanced species, the summed abundance of all species except masu salmon. A Bayesian state-space model is best suited for our analysis because it can account for observation errors while imputing missing values given the long-term trend at each site (*13*, *14*). The model is composed of observation and state models, as described below.

In the observation model, we model observation processes. Fish abundance of either whole community, enhanced species (masu salmon), or unenhanced species at site $s$ in year $t$, $N_{s,t}$, was assumed to follow a Poisson distribution:

$$N_{s,t} \sim Poisson(\lambda_{s,t} A_{s,t}) \tag{5}$$

where $\lambda_{s,t}$ is the expected fish density (individual m$^{-2}$) and $A_{s,t}$ the sampling area (m$^2$). Since fish sampling was conducted after the spring stocking of masu salmon, captured fish may include individuals released in the observation year. We explicitly modeled this observation process to avoid biases in estimating temporal community trends:

$$\lambda_{s,t} = n_{s,t} exp(\epsilon_{s,t}^{obs}) + \psi \beta_s \; Fry_{w(s),t} \tag{6}$$

$n_{s,t}$ is the "true" fish density excluding fish released in the spring, $Fry_{w(s),t}$ the number of salmon fry released (unit: million fish) in spring in watershed $w$ within which site $s$ is located, and $\beta_s$ the site-specific effect of released salmon fry on the observed fish density. The parameter $\beta_s$ was drawn from a normal distribution with the hyper-mean $\mu_\beta$ and hyper-variance $\sigma_\beta^2$. The parameter $\epsilon_{s,t}^{obs}$ is the error term that follows a normal distribution $Normal(0, \sigma_{obs,s}^2)$. The inclusion of this term allows the model to account for site- and year-specific observation errors, which can be caused by ecological and/or artificial factors. When modeling the unenhanced species group, $\psi$ equals zero (otherwise $\psi = 1$) so the model excludes the term $\beta_s \; Fry_{w(s),t}$.

In the state model, we model temporal dynamics of fish density $n_{s,t}$ as follows:

$$ln \; n_{s,t+1} = ln \; r_s + ln \; n_{s,t} + \epsilon_{s,t}^{state} \tag{7}$$

where $ln \; r_s$ is the site-specific rate of change at site $s$, and $\epsilon_{s,t}^{state}$ is the process error that follows a normal distribution as $\epsilon_{s,t}^{state} \sim Normal(0, \sigma_{state,s}^2)$. The site-specific rate of change is random draws from a normal distribution $ln \; r_s \sim Normal(\mu_r, \sigma_r^2)$, assuming that community dynamics across Hokkaido have a shared temporal trend to some degree (the degree of shared trend is controlled by the SD $\sigma_r$). This hierarchical structure allows for improved parameter estimates by partially sharing information across sites (*15*). We used median estimates of fish density $n_{s,t}$ to calculate the temporal CV, mean ($\mu$), and SD ($\sigma$) for each site. We summarized the reconstructed community dynamics in **Figures S7-S9**.

We assessed the predictive performance of our model using the Bayesian p-value (*16*), a value of which takes a range of 0-1 and indicates over- (~0.0), under- (~1.0), or suitable-fitting (~0.5) to the data. Bayesian p-values for our state-space models ranged from 0.48 to 0.51, indicating that our model specification is appropriate.

We used linear regression to quantify the impact of stock enhancement on community dynamics. Although our focus is stock enhancement, each model included climatic and local abiotic variables to account for important environmental differences among sites. Specifically, we developed the following linear regression model taking either the CV, species richness (the number of species present during the observation period), mean, or SD as a response variable $y_s$ with a normal or a Poisson distribution.

$$\begin{cases} ln\ y_s \sim Normal(\mu_{y,s}, \sigma_y^2) & \text{for CV, mean, and SD} \\ y_s \sim Poisson(\lambda_{y,s} exp(\epsilon_{\lambda,s})) & \text{for species richness} \end{cases} \tag{8}$$

where $\sigma_y$ is the SD of residual errors and $exp(\epsilon_{\lambda,s})$ is the error term that accounts for overdispersion ($\epsilon_{\lambda,s} \sim Normal(0, \sigma_\lambda^2)$). The expected means were related to linear predictors as follows:

$$\begin{cases} \mu_{y,s} = \gamma_{0,w(s)} + \sum_k \gamma_k x_{k,s} & \text{for CV, mean, and SD} \\ ln\ \lambda_{y,s} = \gamma_{0,w(s)} + \sum_k \gamma_k x_{k,s} & \text{for species richness} \end{cases} \tag{9}$$

$\gamma_{0,w(s)}$ is the watershed-specific intercept ($w(s)$ refers to site $s$ nested within watershed $w$) and $\gamma_k (k > 0)$ are the regression coefficients of site-level predictors $x_k$. The site-level predictors include upstream watershed area (log-transformed), air temperature, precipitation, and forest land use. Urban and agricultural land use were omitted because of either a limited value range (**Figure S10**) or a strong correlation with forest land use (**Figure S11**). The watershed-specific intercept was related to watershed-level predictors as:

$$\gamma_{0,w} \sim Normal(\mu_{\gamma,w}, \sigma_\gamma^2) \tag{10}$$

$$\mu_{\gamma,w} = \delta_0 + \sum_k \delta_k z_{k,w} \tag{11}$$

$\delta_0$ is the global intercept and $\delta_k (k > 0)$ are the regression coefficients. The watershed-level predictors $z_{k,w}$ include the yearly stocking of masu salmon (fry + juvenile + smolt; averaged for1999-2019) and ocean productivity (chlorophyll *a* concentration; averaged for 2002-2019). Ocean productivity was included because the majority of the observed species use marine habitats at a certain life stage (i.e., diadromous). The parameter $\sigma_\gamma$ accounts for random variation among watersheds that the watershed-level predictors cannot capture. All predictors were standardized (mean = 0, SD = 1) before the analysis.

We fitted the models to the data using JAGS version 4.1.0 through *runjags* package version 2.2.0-2 in R (*17*). We assigned weakly informative priors to parameters (**Table S6**). Three Markov chain Monte Carlo (MCMC) chains were run until parameter estimates converged. Total MCMC interations ranged from $3.5 \times 10^4$ to $4.5 \times 10^4$ for the state-space models and was $1.5 \times 10^4$ for the regression models. The first 5000 iterations were discarded as burn-in, and MCMC samples were saved every 40 (state-space) and 20 iterations (regression) to reduce autocorrelation. Convergence was assessed by examining whether the $\hat{R}$ indicator of each parameter approached $< 1.1$ (*15*). Data manipulation and analysis were performed in R version 4.1.0 (*18*). Parameter estimates were summarized in **Table S7-S10**.

### Supplementary text

#### Sensitivity analysis

We performed a sensitivity analysis of the community simulation to identify key simulation parameters that strongly affect the relationships between community dynamics (temporal mean and SD of density) and stock enhancement. We generated 500 sets of parameter combinations by randomly drawing values of seven simulation parameters from uniform distributions (**Table S1**). For each parameter combination, we simulated dynamics of 100 independent communities with varying numbers of stock enhancement $R$ (drawn randomly from $Unif(0, 500)$). This yields a total of $5 \times 10^4$ simulation replicates. In each simulation replicate, we ran 1600 time steps of community dynamics and obtained temporal means and SDs of the whole community ($\sum_i^S N_i$), enhanced species ($N_1$), and unenhanced species ($\sum_{i,i \neq 1}^S N_i$) using the last 1000 time steps. The first 600 time steps were discarded as initialization and burn-in periods.

For each parameter combination, we estimated Spearman's rank correlation $\xi$ between the number of releases $R$ and community dynamics, which indicate the effect of stock enhancement under a given ecological context. To examine influences of simulation parameters on $\xi$ (**Table S1**), we developed the following regression model taking $\xi$ as a response variable:

$$\xi_n \sim Normal(\mu_n, \sigma^2)$$
$$\mu_n = \zeta_0 + \zeta_1 S_n + \zeta_2 r_{1,n} + \zeta_3 r_{max,n} + \zeta_4 \bar{\alpha}_n + \zeta_5 K_n + \zeta_6 \sigma_{\epsilon,n} + \zeta_7 f_{R,n}$$

where $\xi_n$ is Spearman's rank correlation for parameter combination $n$, and $\zeta_k$ ($k = 0 - 7$) are the intercept ($\zeta_0$) and regression coefficients ($\zeta_{1-7}$). Explanatory variables were standardized (mean = 0, SD = 1) to compare regression coefficients.

We found that regression coefficients of the following parameters ($\zeta$) never exceeded a value of 0.10: the number of species $S$, maximum intrinsic growth rate of unenhanced species $r_{max}$, environmental stochasticity $\sigma_\epsilon$, and relative fitness $f_R$. Therefore, these parameters have little influence on the qualitative relationship between the number of releases and community dynamics. In the main simulation, we fixed values of $S$, $r_{max}$, and $\sigma_\epsilon$ (**Table S1**). In the meantime, we varied values of $K$, $r_1$, $f_R$, and $\bar{\alpha}$ as they are potentially influential (**Table S1**). Although $f_R$ had little influence on stocking effects (**Table S1**), we varied this parameter given the significant interest in the relative fitness of captive-bred individuals in the wild.

### Tables

#### Table S1 Simulation parameter

Parameter values used in the main and sensitivity analysis. Five hundred parameter values were drawn randomly from uniform distributions in the sensitivity analysis.

| Parameter | Interpretation | Main | Sensitivity |
|---|---|---|---|
| $S$ | Number of species | 10 | Unif(5, 20) |
| $r_1$ | Intrinsic growth rate of an enhanced species | 0.5, 1.0, 2.0 | Unif(0.5, 2.5) |
| $r_{max}$ | Maximum intrinsic growth rate of unenhanced species | 2 | Unif(0.5, 2.5) |
| $\bar{\alpha}$ | Average strength of interspecific competition | 0.1, 0.5 | Unif(0.05, 0.5) |
| $K$ | Carrying capacity | 100, 500 | Unif(100, 1000) |
| $\sigma_\epsilon$ | Environmental variability | 0.5 | Unif(0.05, 0.5) |
| $f_R$ | Relative fitness of captive-bred individuals | 0.5, 1.0 | Unif(0.5, 1) |

#### Table S2 Sensitivity analysis (whole community)

Sensitivity analysis of the community simulation. Parameter estimates of linear regression models (standard errors in parenthesis) are shown. Response variables are Spearman's rank correlation between stock enhancement and either temporal mean density ($\mu$) or SD ($\sigma$) of whole community. Explanatory variables (i.e., simulation parameters) were standardized (mean = 0, SD = 1) before the analysis.

| | Response variable | |
|---|---|---|
| | Correlation with $\mu$ | Correlation with $\sigma$ |
| Number of species $S$ | −0.014<br>(0.004) | 0.038<br>(0.006) |
| Intrinsic growth rate $r_1$ | −0.071<br>(0.004) | −0.079<br>(0.006) |
| Maximum intrinsic growth rate $r_{max}$ | −0.003<br>(0.004) | −0.010<br>(0.006) |
| Competition coefficient $\bar{\alpha}$ | 0.036<br>(0.004) | 0.019<br>(0.006) |
| Carrying capacity $K$ | 0.162<br>(0.004) | 0.113<br>(0.006) |
| Environmental variability $\sigma_\epsilon$ | 0.006<br>(0.004) | −0.031<br>(0.006) |
| Relative fitness $f_R$ | 0.020<br>(0.004) | 0.017<br>(0.006) |
| Intercept | −0.481<br>(0.004) | −0.311<br>(0.006) |

#### Table S3 Sensitivity analysis (enhanced species)

Sensitivity analysis of the community simulation. Parameter estimates of linear regression models (standard errors in parenthesis) are shown. Response variables are Spearman's rank correlation between stock enhancement and either temporal mean density ($\mu$) or SD ($\sigma$) of enhanced species. Explanatory variables (i.e., simulation parameters) were standardized (mean = 0, SD = 1) before the analysis.

| | Response variable | |
|---|---|---|
| | Correlation with $\mu$ | Correlation with $\sigma$ |
| Number of species $S$ | 0.060 | 0.041 |
| | (0.009) | (0.008) |
| Intrinsic growth rate $r_1$ | −0.188 | −0.158 |
| | (0.009) | (0.008) |
| Maximum intrinsic growth rate $r_{max}$ | −0.004 | 0.011 |
| | (0.009) | (0.008) |
| Competition coefficient $\bar{\alpha}$ | 0.169 | 0.130 |
| | (0.009) | (0.008) |
| Carrying capacity $K$ | 0.237 | 0.244 |
| | (0.009) | (0.008) |
| Environmental variability $\sigma_\epsilon$ | −0.005 | 0.004 |
| | (0.009) | (0.008) |
| Relative fitness $f_R$ | 0.038 | 0.067 |
| | (0.009) | (0.008) |
| Intercept | −0.223 | −0.336 |
| | (0.009) | (0.008) |

#### Table S4 Sensitivity analysis (unenhanced species)

Sensitivity analysis of the community simulation. Parameter estimates of linear regression models (standard errors in parenthesis) are shown. Response variables are Spearman's rank correlation between stock enhancement and either temporal mean density ($\mu$) or SD ($\sigma$) of unenhanced species. Explanatory variables (i.e., simulation parameters) were standardized (mean = 0, SD = 1) before the analysis.

| | Response variable | |
|---|---|---|
| | Correlation with $\mu$ | Correlation with $\sigma$ |
| Number of species $S$ | −0.051<br>(0.005) | −0.002<br>(0.005) |
| Intrinsic growth rate $r_1$ | 0.016<br>(0.005) | 0.007<br>(0.005) |
| Maximum intrinsic growth rate $r_{max}$ | −0.005<br>(0.005) | −0.014<br>(0.005) |
| Competition coefficient $\bar{\alpha}$ | −0.020<br>(0.005) | −0.023<br>(0.005) |
| Carrying capacity $K$ | 0.107<br>(0.005) | 0.061<br>(0.005) |
| Environmental variability $\sigma_\epsilon$ | 0.004<br>(0.005) | −0.018<br>(0.005) |
| Relative fitness $f_R$ | −0.003<br>(0.005) | −0.007<br>(0.005) |
| Intercept | −0.393<br>(0.005) | −0.293<br>(0.005) |

#### Table S5 Observed species

Fish species found in the study watersheds.

| Taxon | Number of sites observed |
|---|---|
| *Barbatula oreas* | 51 |
| *Cottus* spp. | 66 |
| *Gasterosteus* spp. | 7 |
| *Gymnogobius* spp. | 39 |
| *Lethenteron* spp. | 24 |
| *Luciogobius guttatus* | 7 |
| *Misgurnus anguillicaudatus* | 10 |
| *Oncorhynchus masou masou* | 97 |
| *Oncorhynchus mykiss* | 31 |
| *Parahucho perryi* | 1 |
| *Plecoglossus altivelis altivelis* | 32 |
| *Pseudaspius* spp. | 46 |
| *Pseudorasbora* spp. | 2 |
| *Pungitius* spp. | 13 |
| *Rhinogobius* spp. | 30 |
| *Rhynchocypris lagowskii steindachneri* | 1 |
| *Rhynchocypris percnura sachalinensis* | 1 |
| *Salvelinus leucomaenis leucomaenis* | 94 |
| *Salvelinus malma krascheninnikovi* | 5 |
| *Tridentiger brevispinis* | 4 |

#### Table S6 Priors

Prior distributions used in the Bayesian models.

| Model | Parameter | Prior |
|---|---|---|
| State space | $ln\ n_{s,1}$ | Normal(0, 10) |
| | $\mu_r$ | Normal(0, 10) |
| | $\mu_\beta$ | Normal(0, 10) |
| | $\sigma_r$ | half-Student's t(0, 2.5, 3) |
| | $\sigma_\beta$ | half-Student's t(0, 2.5, 3) |
| | $\sigma_{state,s}$ | half-Student's t(0, 2.5, 3) |
| | $\sigma_{obs,s}$ | half-Student's t(0, 2.5, 3) |
| Regression | $\gamma_k$ | Normal(0, 10) |
| | $\delta_k$ | Normal(0, 10) |
| | $\sigma_y$ | half-Student's t(0, 2.5, 3) |
| | $\sigma_\gamma$ | half-Student's t(0, 2.5, 3) |

#### Table S7 Parameter estimates of the Bayesian state-space model

Median estimates of the Bayesian state-space model. Numbers in parenthesis indicate 95% credible intervals. Site-specific parameters were excluded due to a large number of parameters.

| Species group | Parameter | Interpretation | Estimate |
|---|---|---|---|
| Whole | $\mu_r$ | Rate of community change | $-0.01$ $(-0.03,\ 0.00)$ |
| | $\mu_\beta$ | Effect of spring stoking | 0.64 (0.30, 1.04) |
| | $\sigma_r$ | SD of the rate of community change | 0.01 (0.00, 0.03) |
| | $\sigma_\beta$ | SD of the effect of spring stocking | 0.71 (0.31, 1.17) |
| Enhanced | $\mu_r$ | Rate of community change | $-0.03$ $(-0.06,\ -0.01)$ |
| | $\mu_\beta$ | Effect of spring stoking | 0.45 (0.25, 0.72) |
| | $\sigma_r$ | SD of the rate of community change | 0.02 (0.00, 0.05) |
| | $\sigma_\beta$ | SD of the effect of spring stocking | 0.45 (0.20, 0.75) |
| Unenhanced | $\mu_r$ | Rate of community change | $-0.00$ $(-0.02,\ 0.02)$ |
| | $\sigma_r$ | SD of the rate of community change | 0.01 (0.00, 0.03) |

#### Table S8 Parameter estimates for the regression model (whole community)

Median estimates and standard errors (SEs) of regression coefficients are shown. Pr(> 0) and Pr(< 0) represent the proportion of positive and negative estimates in MCMC samples, respectively (i.e., posterior probability).

| Response | Parameter | Effect | Estimate | SE | Pr(> 0) | Pr(< 0) |
|---|---|---|---|---|---|---|
| CV | $\delta_0$ | Intercept | −1.27 | 0.07 | 0.00 | 1.00 |
| | $\delta_1$ | Stock enhancement | 0.21 | 0.08 | 0.99 | 0.01 |
| | $\delta_2$ | Ocean productivity | 0.11 | 0.10 | 0.87 | 0.13 |
| | $\gamma_1$ | Watershed area | 0.12 | 0.07 | 0.98 | 0.02 |
| | $\gamma_2$ | Air temperature | −0.04 | 0.08 | 0.32 | 0.68 |
| | $\gamma_3$ | Precipitation | −0.02 | 0.08 | 0.39 | 0.61 |
| | $\gamma_4$ | Forest fraction | 0.23 | 0.11 | 0.98 | 0.02 |
| Species richness | $\delta_0$ | Intercept | 1.74 | 0.05 | 1.00 | 0.00 |
| | $\delta_1$ | Stock enhancement | −0.11 | 0.05 | 0.02 | 0.98 |
| | $\delta_2$ | Ocean productivity | −0.15 | 0.07 | 0.02 | 0.98 |
| | $\gamma_1$ | Watershed area | 0.09 | 0.05 | 0.96 | 0.04 |
| | $\gamma_2$ | Air temperature | 0.07 | 0.06 | 0.88 | 0.12 |
| | $\gamma_3$ | Precipitation | −0.06 | 0.06 | 0.13 | 0.87 |
| | $\gamma_4$ | Forest fraction | −0.16 | 0.07 | 0.02 | 0.98 |
| Mean $\mu$ | $\delta_0$ | Intercept | −1.08 | 0.10 | 0.00 | 1.00 |
| | $\delta_1$ | Stock enhancement | −0.21 | 0.11 | 0.03 | 0.97 |
| | $\delta_2$ | Ocean productivity | −0.13 | 0.14 | 0.18 | 0.82 |
| | $\gamma_1$ | Watershed area | −0.23 | 0.07 | 0.00 | 1.00 |
| | $\gamma_2$ | Air temperature | 0.34 | 0.11 | 1.00 | 0.00 |
| | $\gamma_3$ | Precipitation | −0.12 | 0.11 | 0.12 | 0.88 |
| | $\gamma_4$ | Forest fraction | 0.14 | 0.15 | 0.85 | 0.15 |
| SD $\sigma$ | $\delta_0$ | Intercept | −2.33 | 0.10 | 0.00 | 1.00 |
| | $\delta_1$ | Stock enhancement | −0.01 | 0.12 | 0.48 | 0.52 |
| | $\delta_2$ | Ocean productivity | −0.05 | 0.15 | 0.37 | 0.63 |
| | $\gamma_1$ | Watershed area | −0.10 | 0.09 | 0.13 | 0.87 |
| | $\gamma_2$ | Air temperature | 0.26 | 0.13 | 0.99 | 0.01 |
| | $\gamma_3$ | Precipitation | −0.10 | 0.13 | 0.19 | 0.81 |
| | $\gamma_4$ | Forest fraction | 0.34 | 0.17 | 0.99 | 0.01 |

#### Table S9 Parameter estimates for the regression model (masu salmon)

Median estimates and standard errors (SEs) of regression coefficients are shown. Pr(> 0) and Pr(< 0) represent the proportion of positive and negative estimates in MCMC samples, respectively (i.e., posterior probability).

| Response | Parameter | Effect | Estimate | SE | Pr(> 0) | Pr(< 0) |
|---|---|---|---|---|---|---|
| Mean $\mu$ | $\delta_0$ | Intercept | −2.01 | 0.21 | 0.00 | 1.00 |
| | $\delta_1$ | Stock enhancement | −0.08 | 0.23 | 0.37 | 0.63 |
| | $\delta_2$ | Ocean productivity | 0.09 | 0.29 | 0.63 | 0.37 |
| | $\gamma_1$ | Watershed area | −0.24 | 0.12 | 0.03 | 0.97 |
| | $\gamma_2$ | Air temperature | 0.26 | 0.21 | 0.89 | 0.11 |
| | $\gamma_3$ | Precipitation | −0.24 | 0.21 | 0.11 | 0.89 |
| | $\gamma_4$ | Forest fraction | 0.36 | 0.28 | 0.91 | 0.09 |
| SD $\sigma$ | $\delta_0$ | Intercept | −2.70 | 0.16 | 0.00 | 1.00 |
| | $\delta_1$ | Stock enhancement | 0.01 | 0.18 | 0.52 | 0.48 |
| | $\delta_2$ | Ocean productivity | −0.10 | 0.23 | 0.32 | 0.68 |
| | $\gamma_1$ | Watershed area | −0.13 | 0.13 | 0.14 | 0.86 |
| | $\gamma_2$ | Air temperature | 0.05 | 0.18 | 0.62 | 0.38 |
| | $\gamma_3$ | Precipitation | −0.03 | 0.17 | 0.42 | 0.58 |
| | $\gamma_4$ | Forest fraction | 0.36 | 0.25 | 0.93 | 0.07 |

#### Table S10 Parameter estimates for the regression model (unenhanced species)

Median estimates and standard errors (SEs) of regression coefficients are shown. Pr(> 0) and Pr(< 0) represent the proportion of positive and negative estimates in MCMC samples, respectively (i.e., posterior probability).

| Response | Parameter | Effect | Estimate | SE | Pr(> 0) | Pr(< 0) |
|---|---|---|---|---|---|---|
| Mean $\mu$ | $\delta_0$ | Intercept | −2.12 | 0.13 | 0.00 | 1.00 |
| | $\delta_1$ | Stock enhancement | −0.21 | 0.14 | 0.06 | 0.94 |
| | $\delta_2$ | Ocean productivity | −0.26 | 0.18 | 0.08 | 0.92 |
| | $\gamma_1$ | Watershed area | −0.42 | 0.10 | 0.00 | 1.00 |
| | $\gamma_2$ | Air temperature | 0.46 | 0.14 | 1.00 | 0.00 |
| | $\gamma_3$ | Precipitation | −0.06 | 0.13 | 0.31 | 0.69 |
| | $\gamma_4$ | Forest fraction | −0.08 | 0.19 | 0.35 | 0.65 |
| SD $\sigma$ | $\delta_0$ | Intercept | −3.48 | 0.15 | 0.00 | 1.00 |
| | $\delta_1$ | Stock enhancement | −0.07 | 0.16 | 0.33 | 0.67 |
| | $\delta_2$ | Ocean productivity | 0.01 | 0.20 | 0.52 | 0.48 |
| | $\gamma_1$ | Watershed area | −0.36 | 0.12 | 0.00 | 1.00 |
| | $\gamma_2$ | Air temperature | 0.48 | 0.17 | 1.00 | 0.00 |
| | $\gamma_3$ | Precipitation | −0.25 | 0.17 | 0.06 | 0.94 |
| | $\gamma_4$ | Forest fraction | 0.14 | 0.22 | 0.74 | 0.26 |

### Figures

#### Figure S1 Theoretical prediction ($r_1 = 0.5$, $K = 100$)

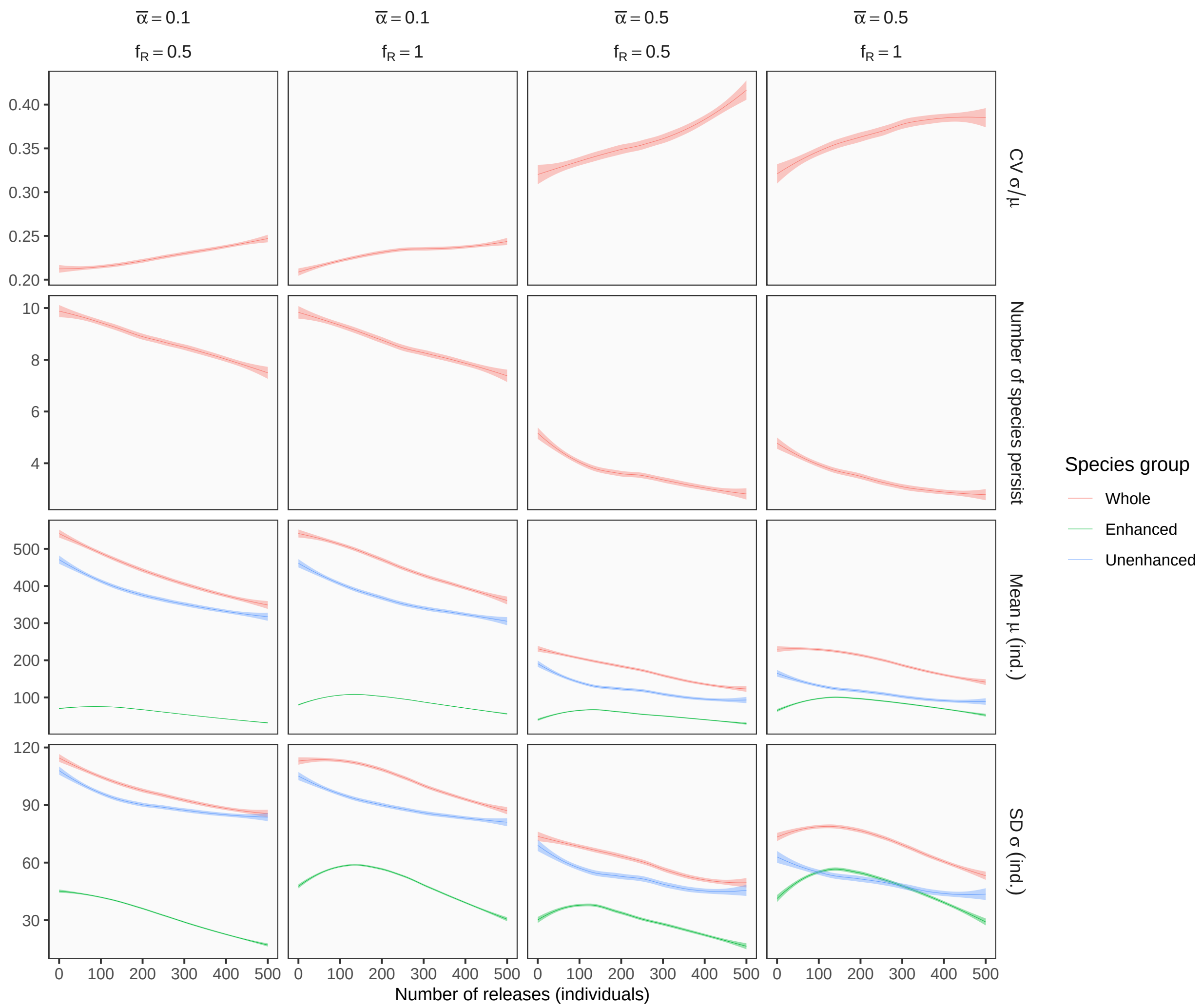

Theoretical predictions for the stocking effect on community dynamics. Rows represent different response variables, and columns show distinct simulation scenarios with different strength of interspecific competition ($\bar{\alpha}$) and relative fitness of captive-bred individuals ($f_R$). Other parameters are: number of species $S = 10$; intrinsic growth rate of an enhanced species $r_1 = 0.5$; maximum intrinsic growth rate of unenhanced species $r_{max} = 2$; environmental stochasticity $\sigma_\epsilon = 0.5$; carrying capacity $K = 100$.

#### Figure S2 Theoretical prediction ($r_1 = 1$, $K = 100$)

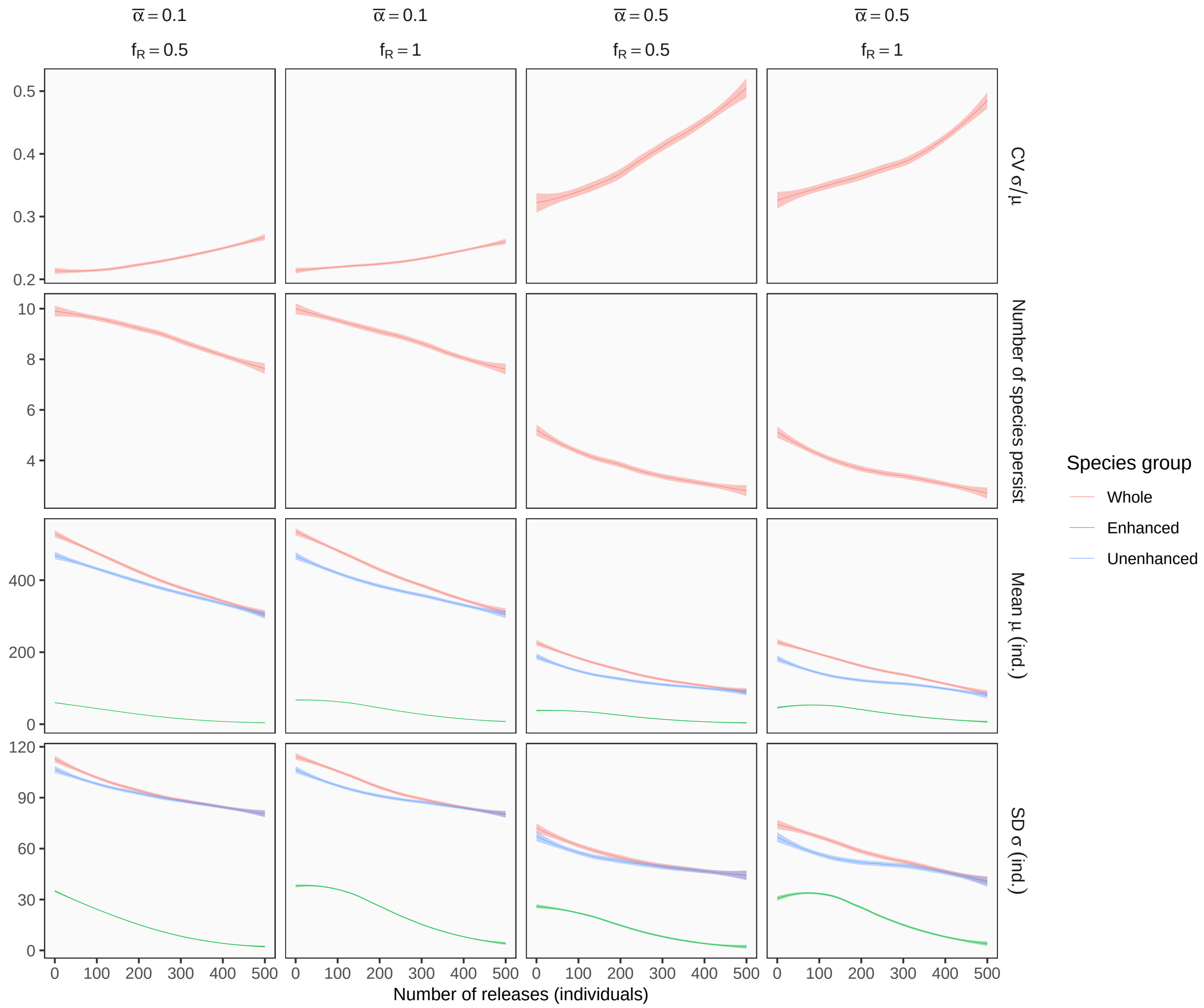

Theoretical predictions for the stocking effect on community dynamics. Rows represent different response variables, and columns show distinct simulation scenarios with different strength of interspecific competition ($\bar{\alpha}$) and relative fitness of captive-bred individuals ($f_R$). Other parameters are: number of species $S = 10$; intrinsic growth rate of an enhanced species $r_1 = 1$; maximum intrinsic growth rate of unenhanced species $r_{max} = 2$; environmental stochasticity $\sigma_\epsilon = 0.5$; carrying capacity $K = 100$.

#### Figure S3 Theoretical prediction ($r_1 = 2$, $K = 100$)

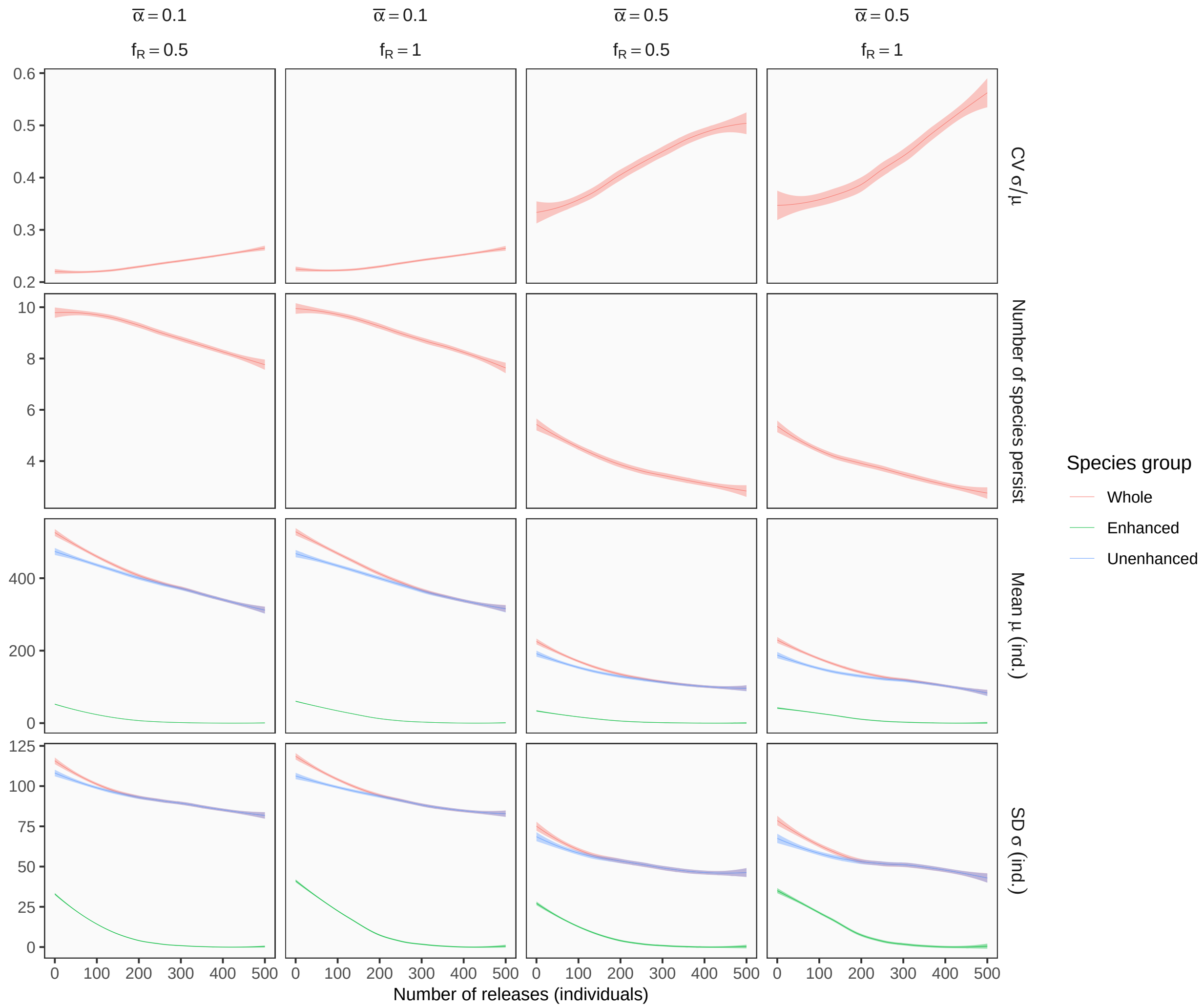

Theoretical predictions for the stocking effect on community dynamics. Rows represent different response variables, and columns show distinct simulation scenarios with different strength of interspecific competition ($\bar{\alpha}$) and relative fitness of captive-bred individuals ($f_R$). Other parameters are: number of species $S = 10$; intrinsic growth rate of an enhanced species $r_1 = 2$; maximum intrinsic growth rate of unenhanced species $r_{max} = 2$; environmental stochasticity $\sigma_\epsilon = 0.5$; carrying capacity $K = 100$.

#### Figure S4 Theoretical prediction ($r_1 = 0.5$, $K = 500$)

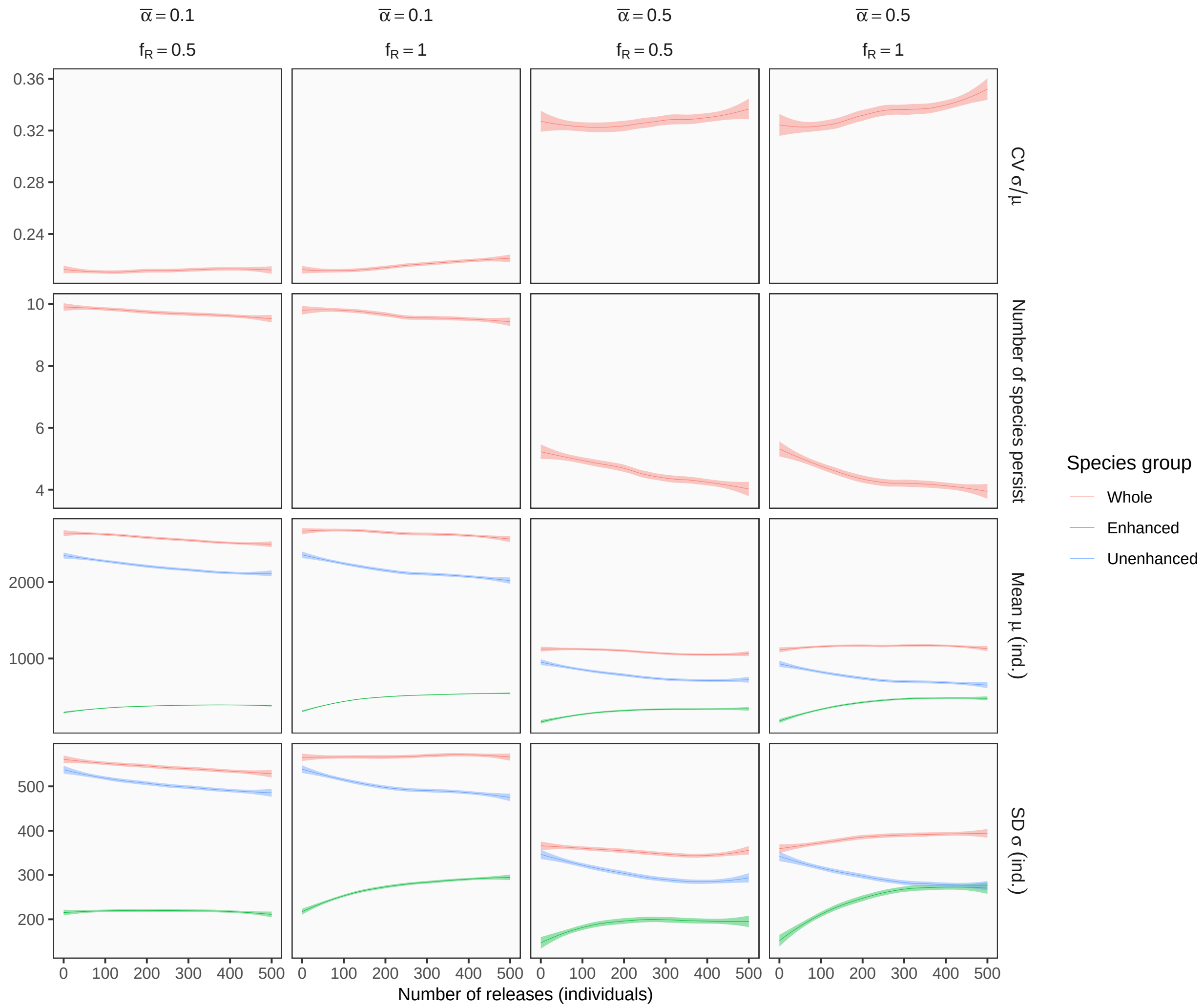

Theoretical predictions for the stocking effect on community dynamics. Rows represent different response variables, and columns show distinct simulation scenarios with different strength of interspecific competition ($\bar{\alpha}$) and relative fitness of captive-bred individuals ($f_R$). Other parameters are: number of species $S = 10$; intrinsic growth rate of an enhanced species $r_1 = 0.5$; maximum intrinsic growth rate of unenhanced species $r_{max} = 2$; environmental stochasticity $\sigma_\epsilon = 0.5$; carrying capacity $K = 500$.

#### Figure S5 Theoretical prediction ($r_1 = 1$, $K = 500$)

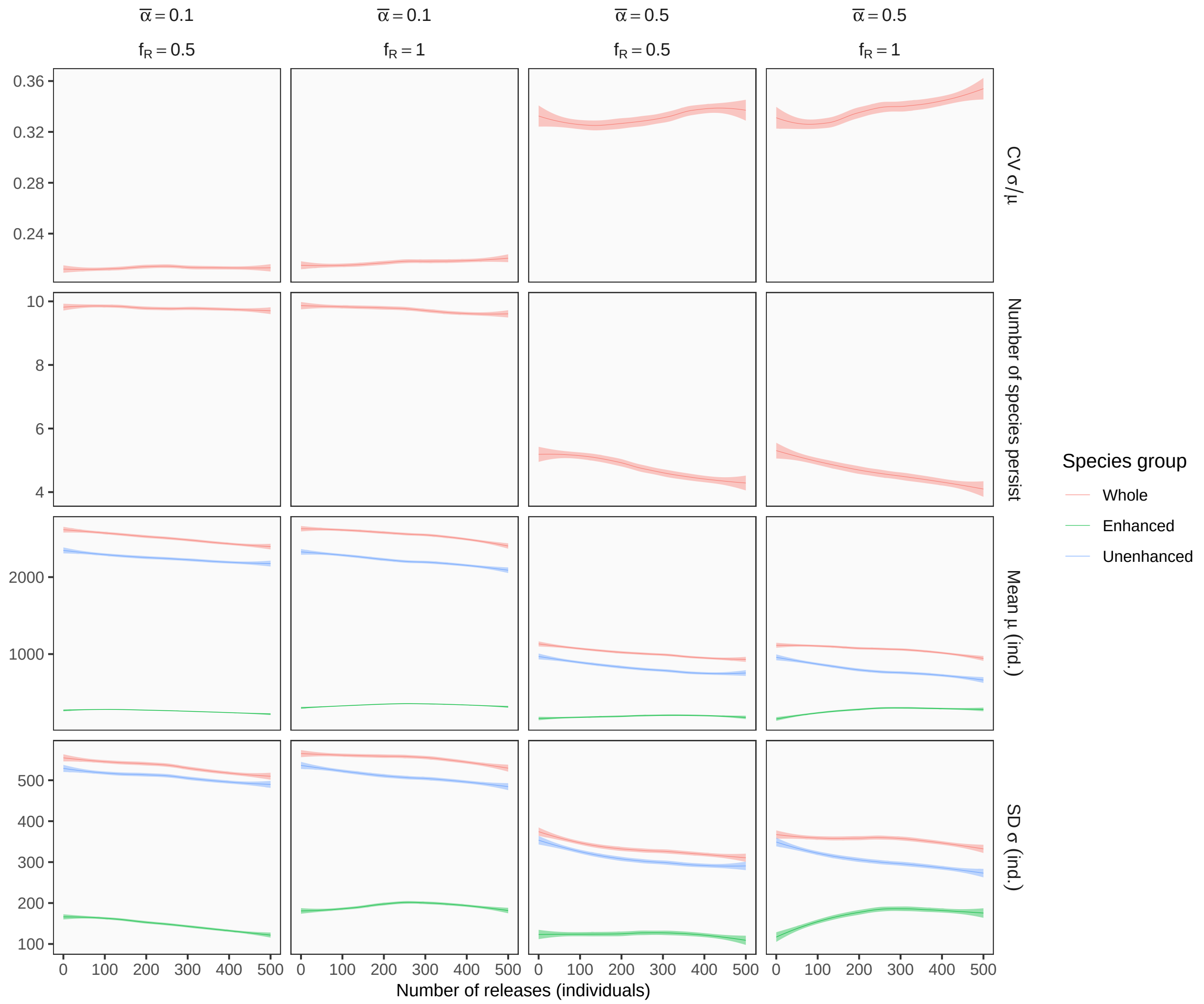

Theoretical predictions for the stocking effect on community dynamics. Rows represent different response variables, and columns show distinct simulation scenarios with different strength of interspecific competition ($\bar{\alpha}$) and relative fitness of captive-bred individuals ($f_R$). Other parameters are: number of species $S = 10$; intrinsic growth rate of an enhanced species $r_1 = 1$; maximum intrinsic growth rate of unenhanced species $r_{max} = 2$; environmental stochasticity $\sigma_\epsilon = 0.5$; carrying capacity $K = 500$.

#### Figure S6 Theoretical prediction ($r_1 = 2$, $K = 500$)

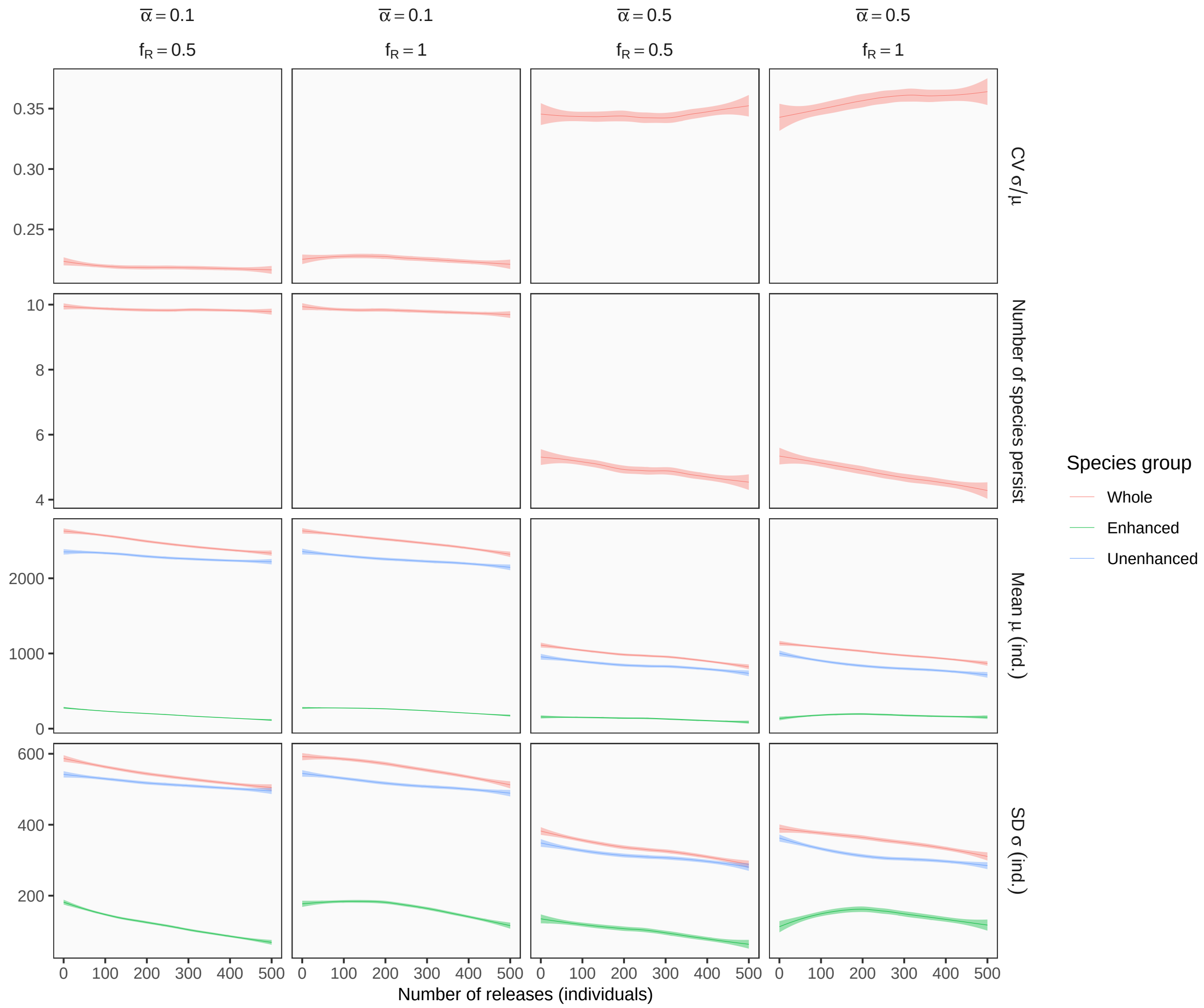

Theoretical predictions for the stocking effect on community dynamics. Rows represent different response variables, and columns show distinct simulation scenarios with different strength of interspecific competition ($\bar{\alpha}$) and relative fitness of captive-bred individuals ($f_R$). Other parameters are: number of species $S = 10$; intrinsic growth rate of an enhanced species $r_1 = 2$; maximum intrinsic growth rate of unenhanced species $r_{max} = 2$; environmental stochasticity $\sigma_\epsilon = 0.5$; carrying capacity $K = 500$.

#### Figure S7 Temporal dynamics of stream fish communities (whole community)

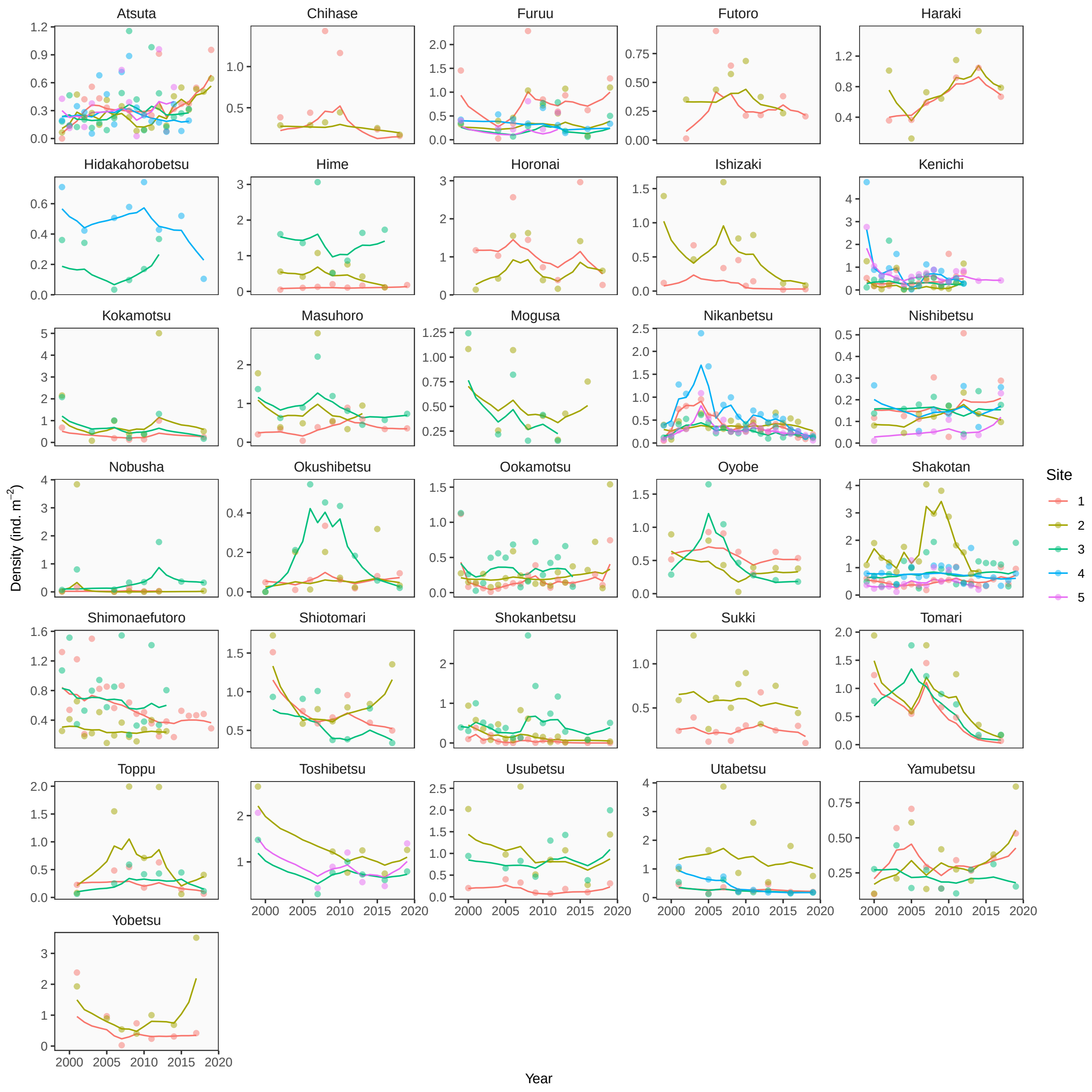

Temporal dynamics of stream fish communities (whole community) in Hokkaido, Japan. Dots represent observed density, and solid lines are the predicted values of the Bayesian state-space model. Panels correspond to individual watersheds, and colors distinguish sampling sites within a watershed.

#### Figure S8 Temporal dynamics of stream fish communities (masu salmon)

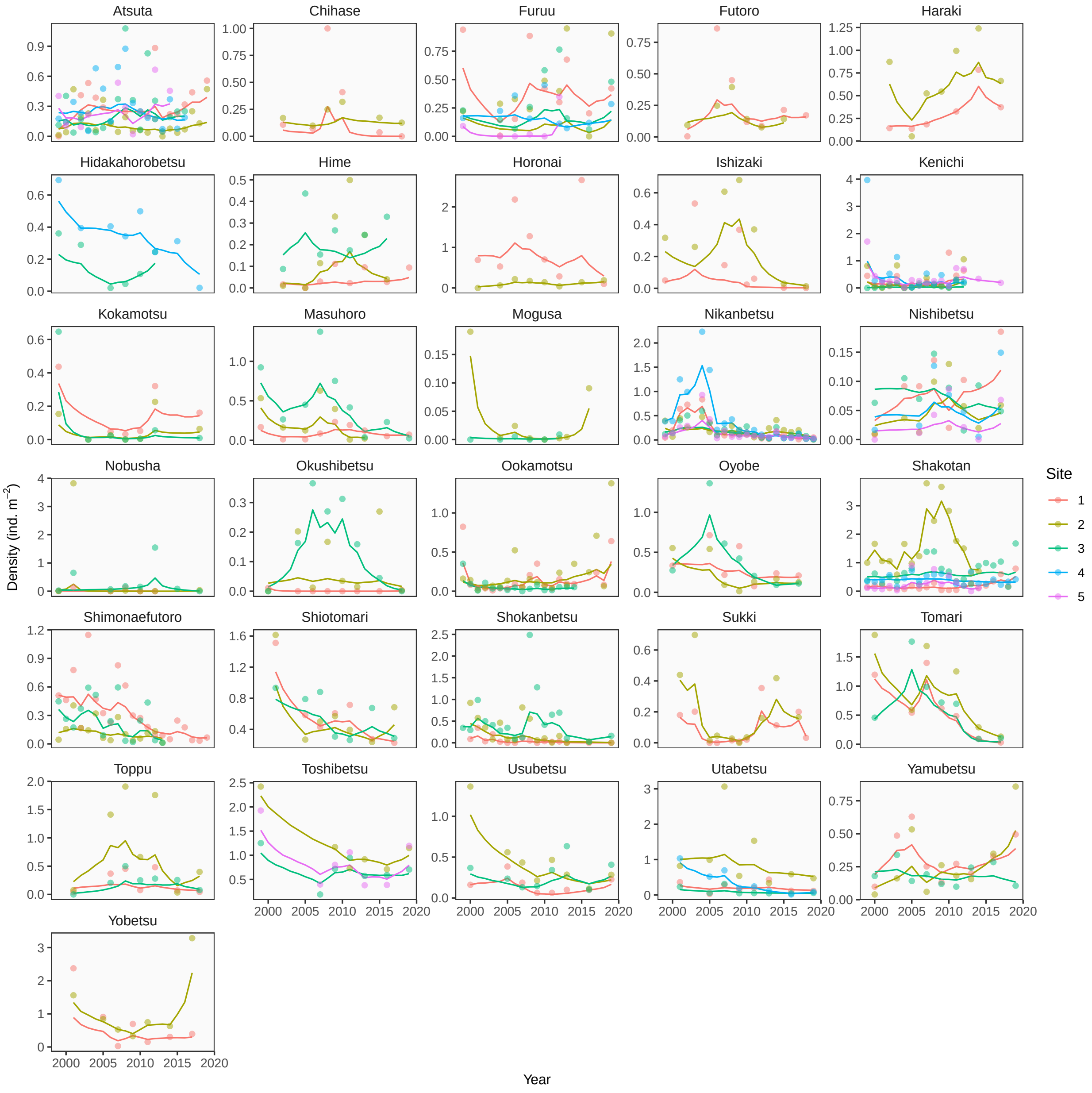

Temporal dynamics of stream fish communities (masu salmon) in Hokkaido, Japan. Dots represent observed density, and solid lines are the predicted values of the Bayesian state-space model. Panels correspond to individual watersheds, and colors distinguish sampling sites within a watershed.

#### Figure S9 Temporal dynamics of stream fish communities (unenhanced species)

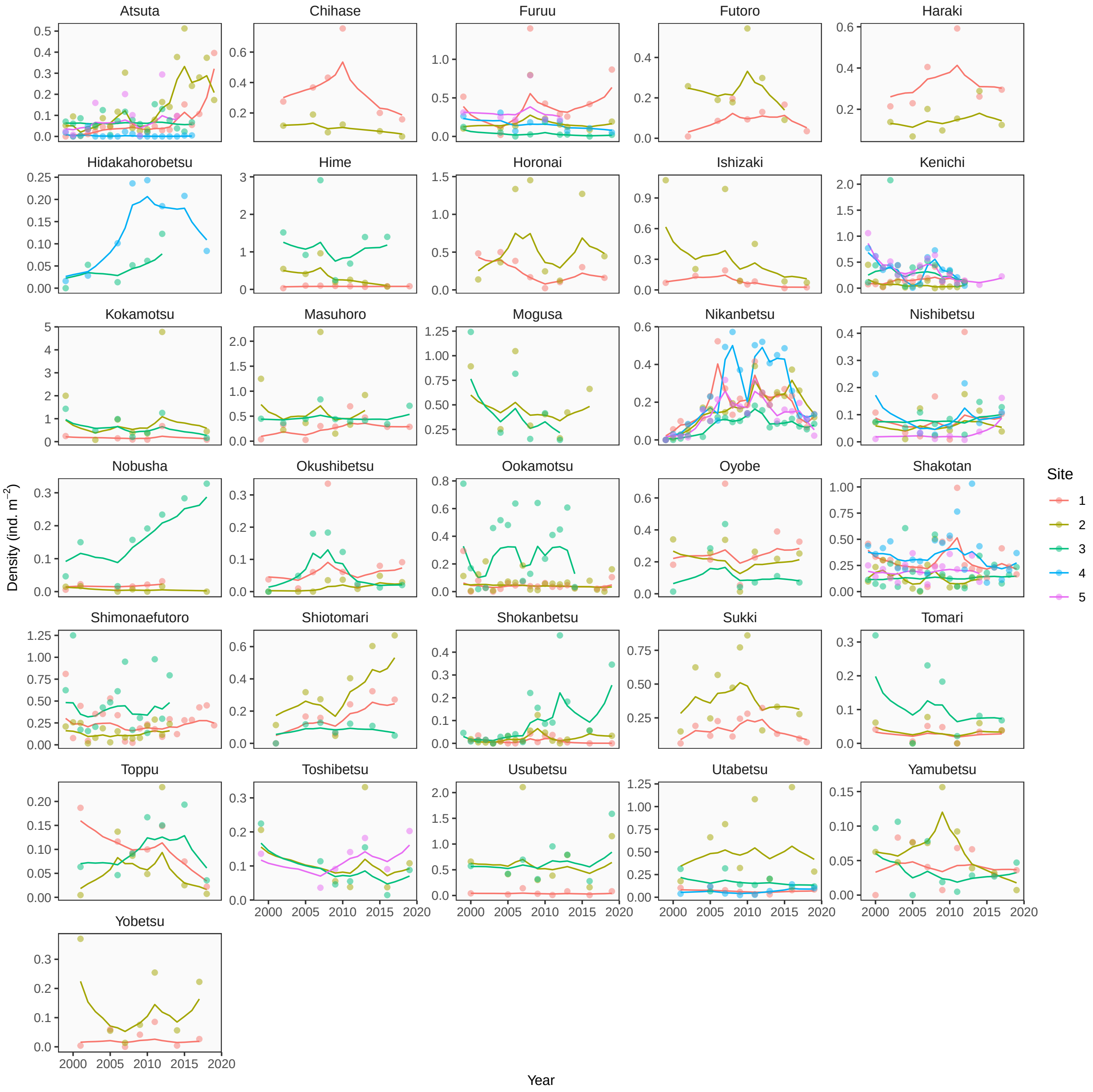

Temporal dynamics of stream fish communities (unenhanced species) in Hokkaido, Japan. Dots represent observed density, and solid lines are the predicted values of the Bayesian state-space model. Panels correspond to individual watersheds, and colors distinguish sampling sites within a watershed.

#### Figure S10 Environmental variables

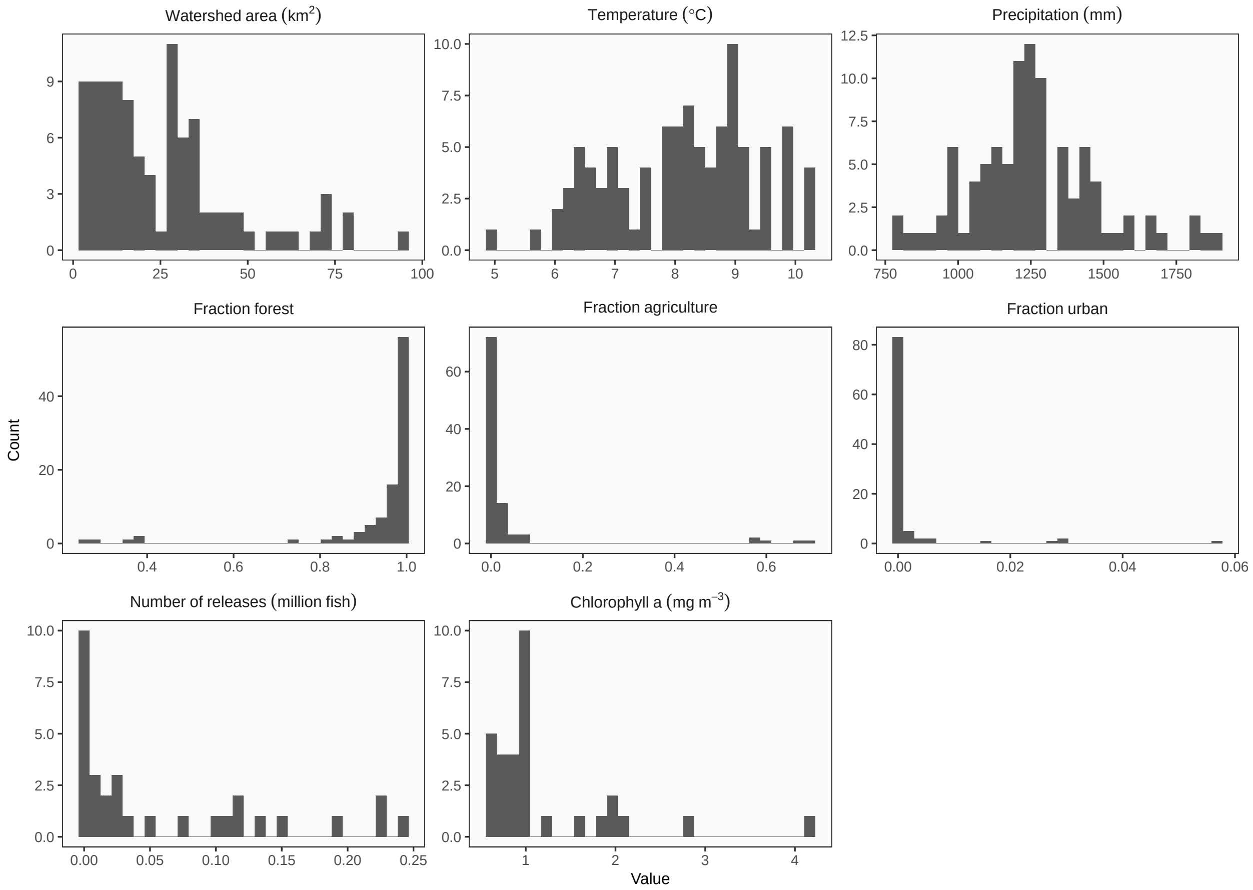

Distribution of environmental variables in the protected watershed. Note that the number of fish released and ocean productivity (chlorophyll *a*) are measured at the watershed level while others are measured at the site level.

#### Figure S11 Correlation plot

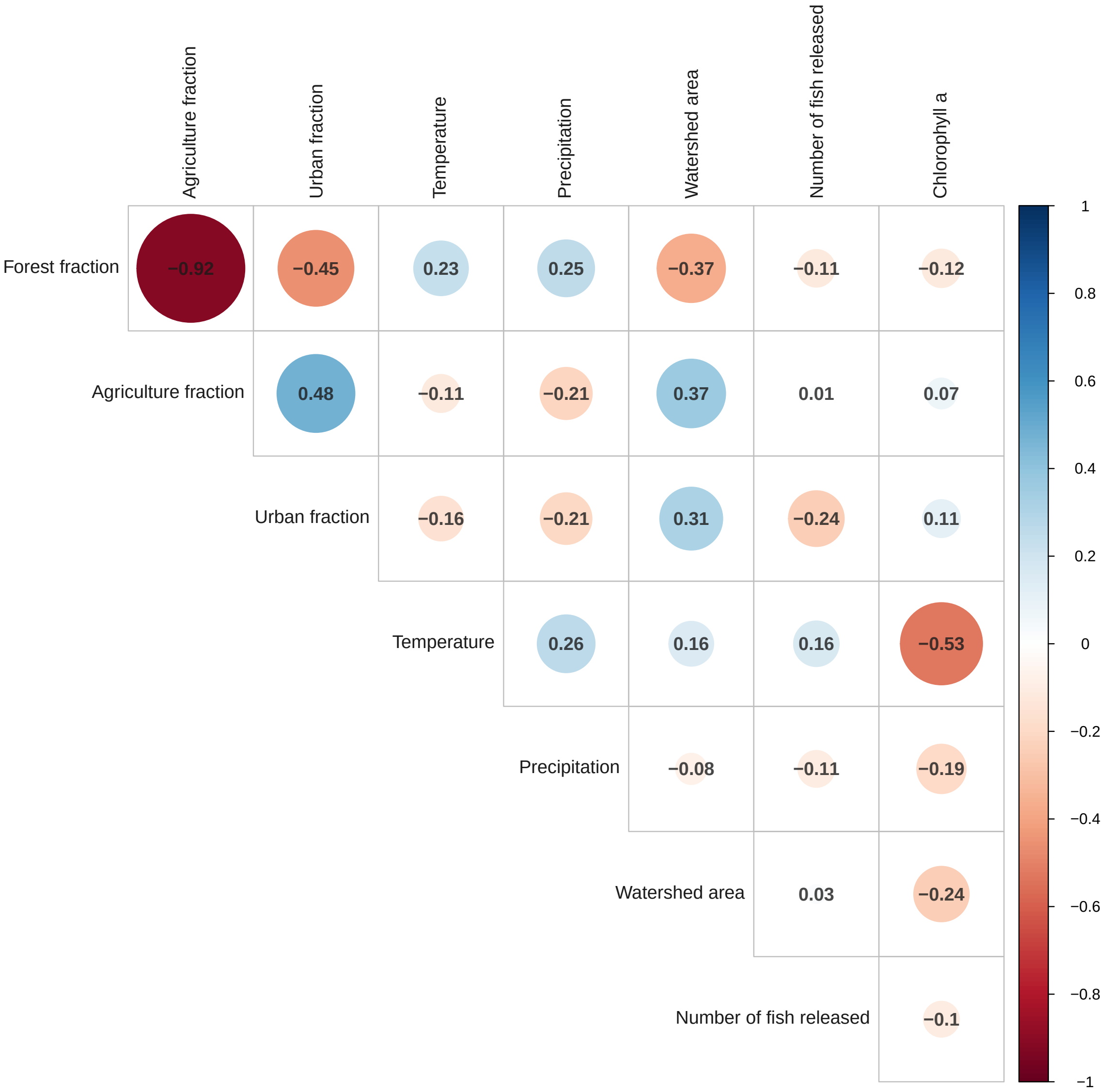

Correlation plot among environmental variables in the protected watersheds. Numbers indicate Spearman's rank correlations.
